# Dynamic protrusions mediate unique crawling motility in Asgard Archaea (Promethearchaeota)

**DOI:** 10.1101/2025.11.30.690169

**Authors:** Philipp Radler, Tobias Viehboeck, Zhen-Hao Luo, Nevena Maslać, Katharina Schmidt, Robert Hauschild, Masaru Nobu, Silvia Bulgheresi, Theresia E.B. Stradal, Klemens Rottner, Hiroyuki Imachi, Michael Sixt, Christa Schleper

## Abstract

Crawling motility is a hallmark of eukaryotic cells and requires a dynamic actin cytoskeleton, regulated adhesion, and spatially organized signalling pathways_1–3_. Asgard archaea (phylum Promethearchaeota) which are considered the closest known prokaryotic relatives of eukaryotes potentially encode these functions within their large set of ‘eukaryotic signature proteins’^4–9^. The few cultivated members show a complex cell morphology, consisting of a central cell body from which several protrusions extend, filled with an actin-based cytoskeleton^10,11^. Here, live cell microscopy of two organisms of the Loki- and Hodarchaea lineages^10,12^ showed that they dynamically and drastically change their cell shape on a minute time scale and grow and retract their extensive protrusions with a speed of 1.5 to 5.3 µm/min, respectively. After adhering to a glass surface, cells employ their protrusions to undergo active crawling motion. In the presence of selected actin inhibitors however, the observed dynamics were arrested, suggesting a central role of actin in these processes. The observed cellular plasticity and motility are unique features among prokaryotes and might have been crucial for the emergence of the first eukaryotic cells that are thought to have formed through the association of a member of the Promethearchaeota and an alphaproteobacterium, the ancestor of mitochondria.

## Introduction

Since their discovery a decade ago, Asgard archaea (now formally classified as the phylum Promethearchaeota^7,11^) have inspired novel models for eukaryogenesis^13,14^. Their genomes encode numerous so-called eukaryotic signature proteins (ESPs), including e.g. ESCRT and ubiquitination machineries, an actin-based cytoskeleton and associated proteins, as well as many small GTPases that were never found to this extent in other prokaryotes^4,5,7^. The members of Promethearchaeota are currently regarded as the closest prokaryotic relatives of eukaryotes and possibly provided their direct ancestor about 2 billion years ago^9,12,15^. While current models of eukaryogenesis differ in their mechanistic details, most of them agree that eukaryotes emerged from a symbiosis or merger between an archaeon of the Promethearchaeota lineage and at least one α-Proteobacterium (or related), which evolved into mitochondria^16^.

The few so-far cultivated cells of the phylum Promethearchaeota^10,11,13,17,18^ have revealed a distinctive cell morphology: a central cell body from which protrusions of varying numbers and lengths extend. Furthermore, the outer membrane was not covered by a cell wall but found to be decorated with proteins, potentially employed for establishing cell–cell contacts or formation of substrate adhesions^10^. The protrusions are filled with a cytoskeleton formed by the conserved actin homologue of all Promethearchaeota, referred to as Lokiactin, which is one of the most highly expressed proteins in *Candidatus* Lokiarchaeum ossiferum ("L. ossiferum") and is both sequence-based and structurally closely related to eukaryotic actins^10,19–21^. Biochemical studies have demonstrated that profilin and gelsolins from Promethearchaeota can bind and modulate eukaryotic actin, further demonstrating the close functional and evolutionary relationship of these proteins^22–24^.

By analogy to eukaryotic pseudopodia, dynamic regulation of Lokiactin filaments could enable protrusion growth and retraction, thereby contributing to morphological plasticity and motility. To directly study if these processes occur in Promethearchaeota, we established anaerobic live-cell imaging and demonstrated that both, L. ossiferum and *Ca.* Margulisarchaeum peptidophilum ("M. peptidophilum") undergo striking morphological rearrangements within minutes. In addition, both Promethearchaeota strains exhibit crawling motility that involves extensive dynamics of their protrusions.

## Results

### Highly dynamic shapes of L. ossiferum cells

We found L. ossiferum to adhere to glass surfaces within two hours of incubation more efficiently than their syntrophic partners of the same culture (Halodesulfovibriae or Spirochaetes, **Fig. S1a**). Most adherent cells (89.6 ± 13.0 %) showed the characteristic morphology of a central cell body with protrusions extending to all sides (**Fig. 1a)**. The number and length of protrusions varied greatly, with an average of five per cell (from 1-10) and a length of 3.5 µm (from 1 to 15 µm) **(Fig. 1b-d)**. Assuming protrusions as a regular tubus of 0.26 µm (0.2 - 0.4 µm) and a perfectly round cell of 1.06 µm in diameter (0.8 - 1.4 µm), we approximated the volume of a single protrusion to be one third of the cell body, but nearly doubling the available surface area. Thus, five protrusions of average length increase the available surface area by 4.5-fold.

**Figure 1:**
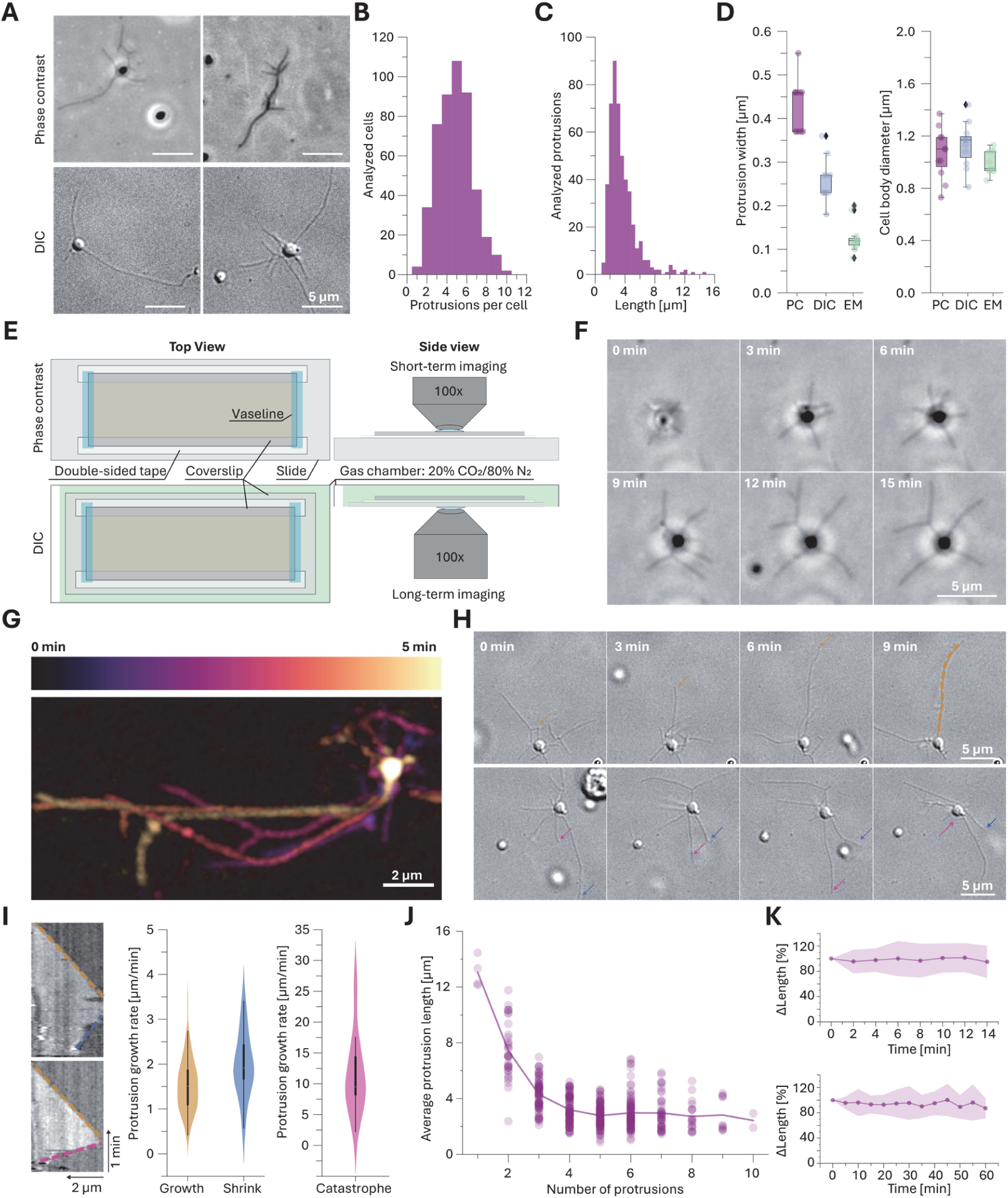
Anaerobic live cell imaging of *Ca.* L. ossiferum reveals plastic cell shape A, Representative micrographs from phase contrast (PC) and differential interference contrast (DIC) microscopy of L. ossiferum with different morphologies. **B/C,** Quantifications of protrusion numbers per cell **(B)** and their lengths **(C)**. **D,** Quantification of the protrusion and cell body diameters using PC, DIC and Scanning Electron Microscopy (SEM). **E,** Scheme of the anaerobic imaging setup with PC (top) and DIC (bottom). Representative experiments are shown in **Supp. Movie S1. F,** Representative PC time lapse micrographs of a L. ossiferum cell adhering to the glass slide. The corresponding experiment is shown in **Supp. Movie S2. G,** A temporal projection of L. ossiferum, illustrating the rearrangement of protrusions over time. **H,** DIC time lapse micrographs of a L. ossiferum cell with growing (**top**) and shrinking (**bottom**) protrusions. The arrows indicate growing (orange), shrinking (blue) and catastrophic (magenta) protrusions. The corresponding experiment is shown in **Supp. Movie S3. I,** Representative kymographs and the corresponding quantification (n = 115 protrusions from 28 experiments of 13 different cultures). **J,** The average protrusion length and the number of protrusions per cell are inversely correlated. **K,** Quantification of the change of total protrusion length of single L. ossiferum cells (n = 46 cells and 220 protrusions).

Apart from the above-described morphology, we also found L. ossiferum with deviating shapes, particularly elongated, stick-like cells that were lacking a central cell body **(Fig 1a, Fig. S1b)**. This morphology constituted the minority of cells (10.4 ± 13.0 %, **Fig. S1c**) and its ratio remained constant throughout the lifetime of a single culture (**Fig. S1d)**, while the total number of attached L. ossiferum cells declined slightly over the course of two weeks (**Fig. S1e-f**). The different cell shapes we observed, suggested either the presence of different cell subtypes or a dynamic re-arrangement of the cell morphology. To differentiate between these possibilities, we established time lapse microscopy by upright phase contrast (PC) and inverted differential interference contrast (DIC) microscopy, while keeping the cells in an anaerobic environment **(Fig 1e, Movie S1)**. Upon contact with the glass surface L. ossiferum cells rapidly spread their protrusions while remaining anchored on the coverslip **(Fig. 1f, Movie S2**). Despite the slow doubling time of the organisms (∼7 days), cells changed their morphology rapidly on a minute time scale. In many cases the growing protrusions branched off to form side arms resulting in two or sometimes more endings per protrusion. Moreover, protrusions grew and retracted constantly, potentially reacting to their environment (**Fig. 1g-h, S1g, Movie S3**). Single protrusions could reach up to 15 µm before they retracted either slowly or in a fast manner. We used kymography to quantify these dynamics and found protrusions to grow at 1.5 ± 0.6 µm/min, and retractions to occur in two distinct categories, either at 2.0 ± 0.7 or at 11.4 ± 6.1 µm/min (**Fig. 1I**). The number and length of protrusions of adhered L. ossiferum cells were inversely correlated, suggesting precise control over the available lipids for protrusion formation (**Fig. 1j**). During the observed rearrangements, the total protrusion length of single cells remained constant, further suggesting a controlled redistribution of the available lipids. (**Fig. 1k**). Finally, we observed significant shape changes of L. ossiferum cell bodies: Merging two cell body-like structures into one (**Fig. S1h**) or switching from a stick-like to a spherical appearance (**Fig. S1i**, **Movie S4**). These morphological changes were not restricted to the cell body alone, as the chromosome, revealed by life staining with NucBlue, followed this dynamic rearrangement **(Fig. S1j, Movie S5)**.

### L. ossiferum migrates on glass using its protrusions

Besides the plastic cell morphology and the rearrangement of protrusions, we found a considerable fraction of L. ossiferum translocating their central cell body (51.4 ± 22.5 %, **Fig S2a**). This movement seemed reminiscent of crawling migration observed in eukaryotes (**Fig. 2a**). The majority of mobile cells migrated randomly over the glass surface, constantly changing their trajectories **(Fig. 2b**). This was reflected in a low confinement ratio, which is a measure of the displacement of the cell body from its starting point **(Fig. 2c)**. Only a small subset of L. ossiferum exhibited persistent motion, as if aimed towards a signal, possibly caused by an attractant emanating from the surroundings (**Movie S6**, 6 of 117 cells). The median speed of migrating L. ossiferum cell bodies was 1.57 ± 0.93 µm/min **(Fig. 2d, S2b-c)**. This is within the range of reported crawling speeds of eukaryotic cells (0.1 – 20 µm/min)^25–27^, but much slower compared to the swimming motility exhibited by Archaea, Bacteria or flagellates (0.6 - 30 mm/min)^28–30^. Moreover, the measured migration velocities of cell bodies were remarkably similar to the observed protrusion growth rates, suggesting a connection between these two. Upon coating the glass with poly-L-lysine, we found a decrease in both the protrusion dynamics and the migration speed (**Fig. S2d-g),** suggesting an intricate interplay between adhesion and detachment from the substrate to allow efficient cell migration^31^.

**Figure 2:**
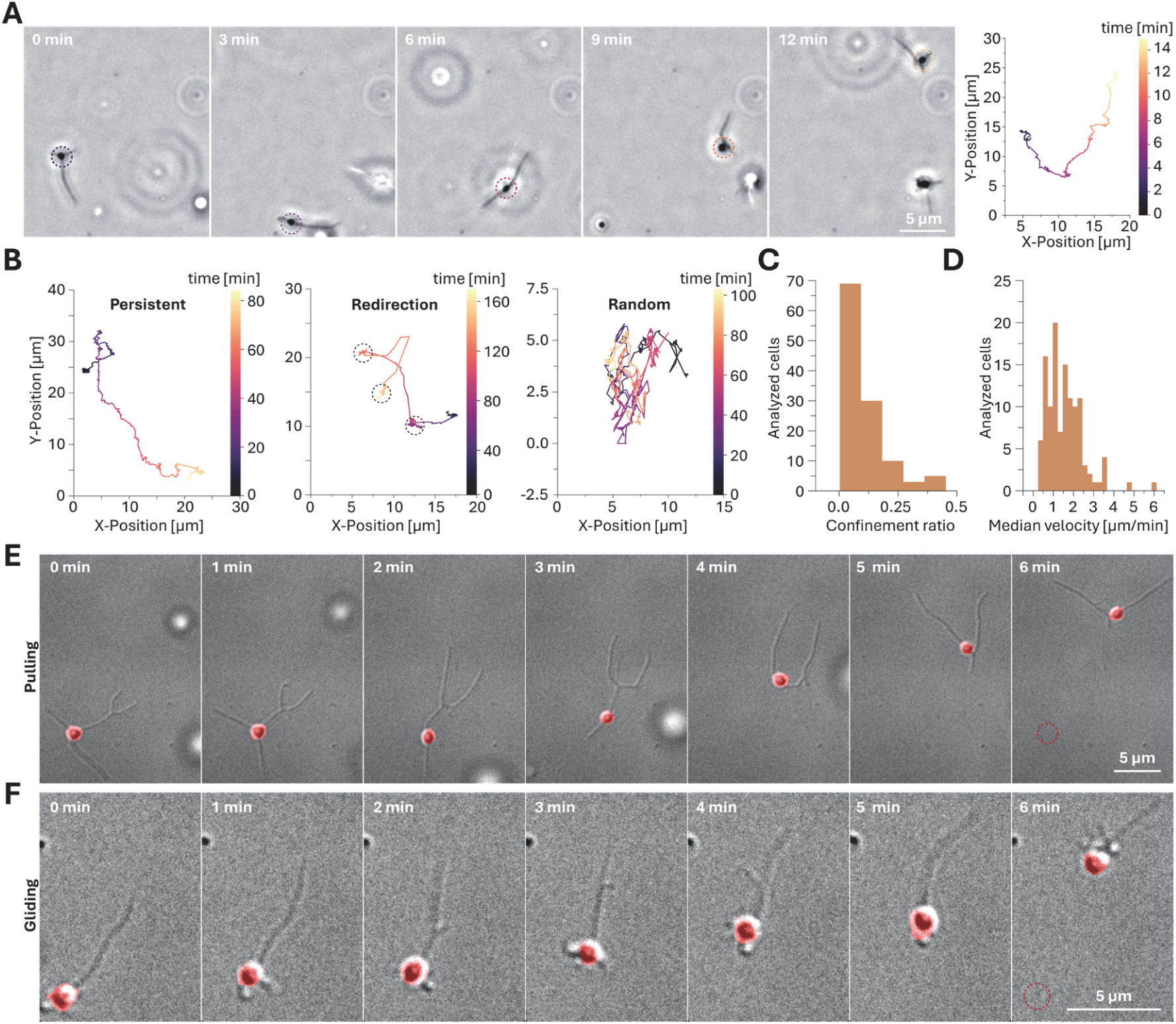
L. ossiferum migrates along glass by protrusion based crawling motility A, Representative PC time lapse micrographs of migrating L. ossiferum and the corresponding trajectory. The dashed circles indicate the position of the cell body at different time points in the track. **B,** Example trajectories of representative L. ossiferum cells moving persistently (left), undergoing several redirections (centre) and moving completely randomly (right). **C/D,** TrackMate quantification of the confinement ratio (**C**) and the median instantaneous velocity **(D)** of L. ossiferum (n=117 cells from 9 different cultures). **E/F,** Representative DIC micrographs of L. ossiferum displacing their cell body via pulling **(E)** and gliding **(F)**. The cell body is shown in red, and the red dashed circle indicates the respective starting point of the cell body. The corresponding experiments are shown in **Supp. Movies S6 and S7**.

We investigated three conceivable mechanical modes of force transduction that can result in crawling motion: pushing, pulling and gliding. Microscopic observations suggest that L. ossiferum employs the latter two: Pulling cells adhered with the tip of leading protrusions and the cell body followed. This mode of mobility required one or more protrusions to retract to move the cell body **(Fig. 2e)**. Gliding motion occurs in the absence of shape changes, because the force transduction happens at a molecular scale, such as treadmilling of actin filaments^1,32–34^. This was seen in particular for cells that exhibited a directionally persistent motion, as the cell bodies followed a single protrusion aimed towards the front (**Fig. 2f, Movie S7**). This type of motion was not interrupted by constant reorientations. To corroborate our microscopic observations, we quantified the alignment between the cell body movement and the direction into which protrusions extend (n = 19 cells). We found the movement direction to be predominantly determined by protrusions in front of the cell body and no evidence for pushing. The alignment was improved when taking the length of protrusions into account (**Fig. S2H, Movie S8**).

### Observed dynamics are dependent on Lokiactin filaments

Next, we investigated if and how far filaments formed by Lokiactin, which are found in almost all protrusions of L. ossiferum (**Fig. S3a**^10,35^), are involved in the observed cell dynamics. In the absence of genetic tools and fluorescent fusion proteins, we first tried to visualize Lokiactin with fluorescent stains developed for eukaryotic actin. SiR Actin or a derivative thereof, FastAct_X, showed very weak fluorescent signal, suggesting a weak affinity to Lokiactin filaments (**Fig. S3b**). Another SiR Actin derivative (“FastAct”) improved staining and signal of L. ossiferum, while retaining high specificity (**Fig. 3a and S3c-d**). While the rates of protrusion growth dropped slightly (0.72 ± 0.49 µm/min, **Fig. 3b**), protrusions still rearranged dynamically (**Fig. 3c, Movie S9**) and actin stain was observed in all dynamic protrusions as well as in the central cell body.

**Figure 3:**
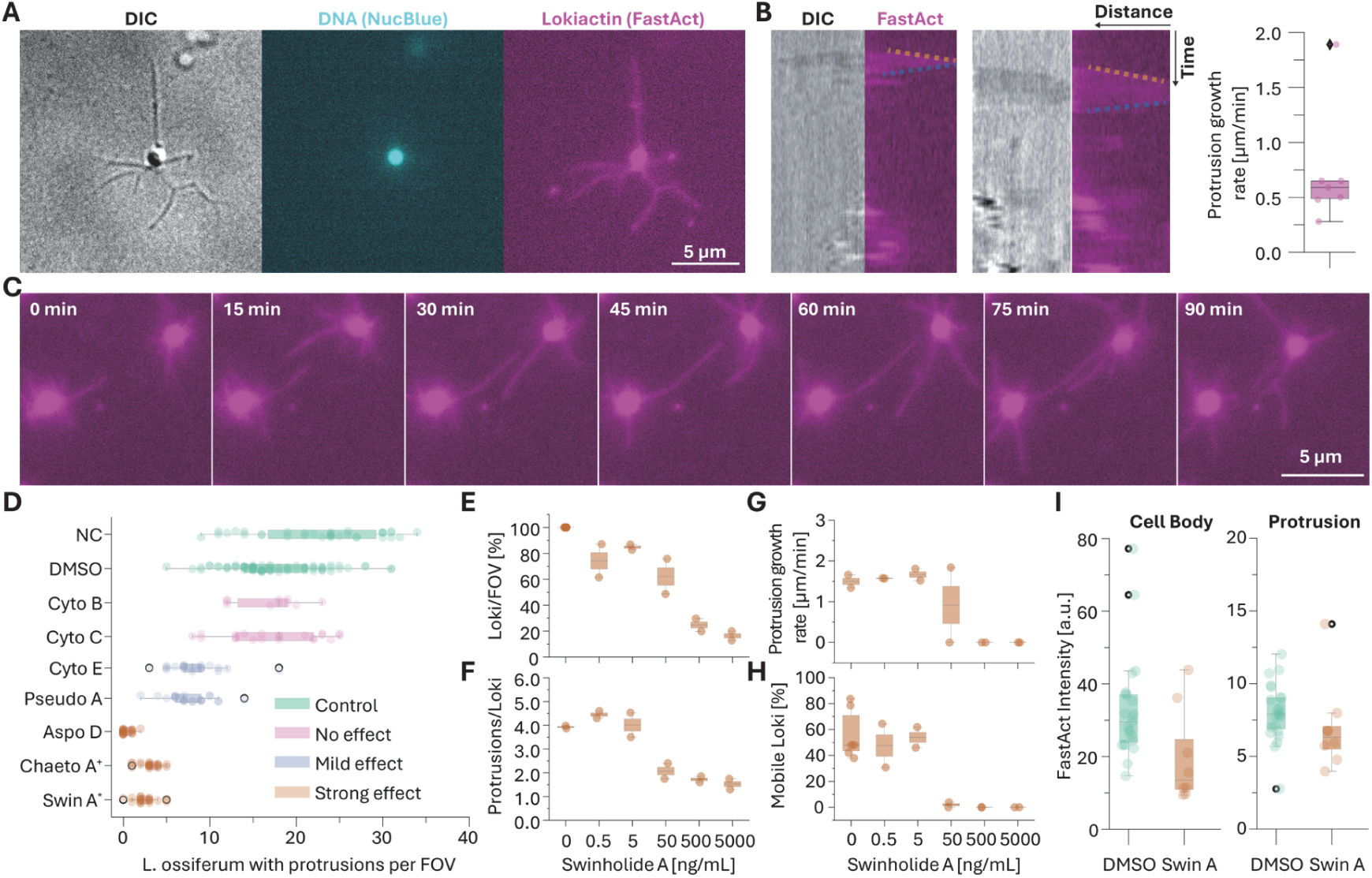
Lokiactin polymerization is essential for protrusion growth and motility A, Representative DIC micrographs of L. ossiferum stained with NucBlue (DNA, cyan) and FastAct (Lokiactin, magenta) **B,** Representative kymographs of growing protrusions in the DIC and FastAct channel together with the corresponding quantification (n = 6 cells). **C,** Example time lapse micrographs of FastAct stained L. ossiferum cells rearranging their protrusions. Corresponding experiments are shown in **Supp. Movie S9**. **D,** Effect of different actin inhibitors on the observed number of L. ossiferum with protrusions per field of view (FOV). The inhibitor concentrations were 25 (^+^), 12.5 or 5 µg/mL (*). **E-H,** Quantification of the number of L. ossiferum with protrusions per FOV **(E)**, the number of protrusions per cell **(F)**, the growth rate of protrusions **(G)** and the percentage of mobile L. ossiferum **(H)** with increasing concentrations of Swinholide A. The titrations were performed with two cultures and we counted 325 L. ossiferum cells with protrusions and quantified kymographs from 43 cells. **I,** Quantification of the FastAct intensity in cell body and protrusions of L. ossiferum treated with 100ng/mL Swin A (n = 23 for DMSO and n = 8 for Swinholide A treated cells).

Next, we wanted to arrest the polymerization dynamics of Lokiactin and tested different inhibitors known to inhibit eukaryotic actin polymerization at high concentrations to identify a compound suitable for L. ossiferum. While some commonly used inhibitors affected the dynamics or morphology only slightly, we found three inhibitors (Aspochalasin D, Chaetoglobosin A and Swinholide A) to significantly reduce the number of L. ossiferum cells adherent to glass **(Fig. 3d, S3e, Supplementary table 1**). We performed a more detailed titration of these inhibitors and found Swinholide A (further referred to as Swin A) to affect L. ossiferum at concentrations as low as 50 ng/mL, within the range of concentrations used in eukaryotic cells^36^ (**Fig. 3e, S3f-g)**. We analysed L. ossiferum cell morphology in detail and found Swin A to decrease the number of protrusions per cell to only one or two remaining appendages (**Fig. 3f**). Moreover, the formation of new protrusions and growth of persisting protrusions was arrested at 50 ng/mL SwinA (**Fig. 3g**). Finally, L. ossiferum lost the ability to migrate and remained stuck in place in the presence of 50 ng/mL Swin A (**Fig. 3h**). Some cells remained attached with a single protrusion that lost internal stabilizing filaments. These protrusions swayed strongly via Brownian motion, which was in stark contrast to the controlled protrusion dynamics observed in the controls (**Movie S10)**. When we combined fluorescence microscopy with Swin A, we found the fluorescent signal in the cell bodies and protrusions to drop slightly, although not disappearing completely (**Fig. 3i, S3h-i**).

### Motility and membrane plasticity are shared characteristics of Promethearchaeota

Next, we were interested in whether the observed dynamics are a common characteristic of Promethearchaeota and performed experiments with M. peptidophilum strain HC1, a representative of the Hodarchaeales lineage^12^. As observed for L. ossiferum, M. peptidophilum cells rearranged their morphology drastically (**Fig. 4a, Movie S11**). In alignment with previous observations^12^, the cell body and protrusions were bigger and longer respectively, than in L. ossiferum (**Fig. 4b-c**). When we measured protrusion dynamics, we found them to grow approximately 3 times faster than protrusions of L. ossiferum (5.3 ± 1.3 vs. 1.5 ± 0.6 µm/min, **Fig. 4d**). M. peptidophilum cells also migrated along the glass randomly, most likely due to a lack of a target (confinement ratio 0.2 ± 0.1 vs. 0.07 ± 0.1, **Fig. 4e-f**). In concordance with the faster protrusion dynamics, we found it to displace its cell body three times faster than L. ossiferum (4.9 ± 1.5 vs. 1.6 ± 0.9 µm/min, **Fig. 4g)**, further suggesting a connection between protrusion and cell migration dynamics. Finally, addition of Swinholide A arrested the rearrangement of cell morphology and migration as observed in L. ossiferum (**Fig 4h, Movie S12**).

**Figure 4:**
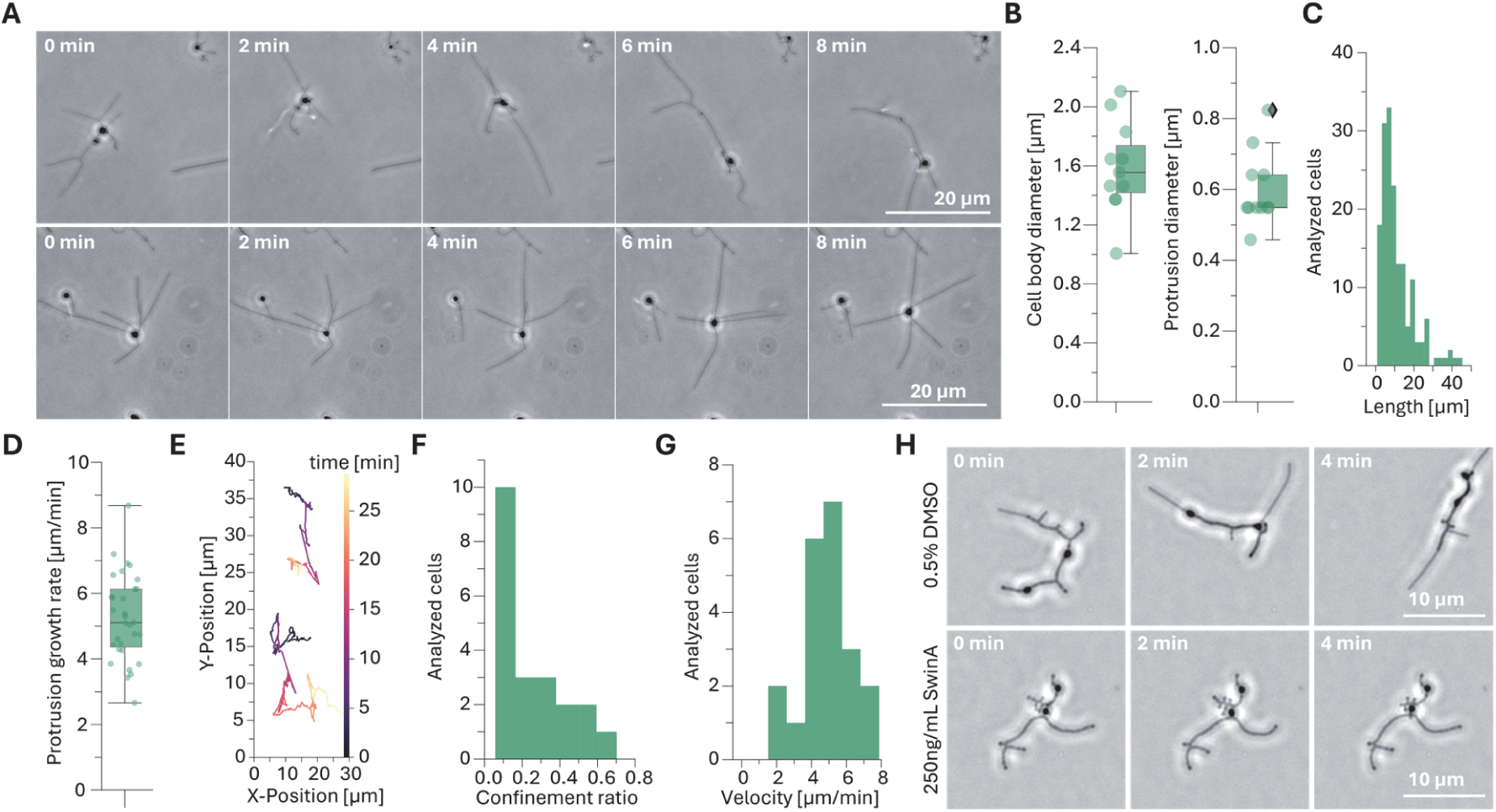
M. peptidophilum displays actin dependent morphological rearrangements A, Representative PC time lapse micrographs of M. peptidophilum rearranging their protrusions and migrating over the glass. The corresponding experiments are shown in **Supp. Movie S11**. **B,** Quantification of the diameter of the cell body and protrusions (n = 11). **C,** Quantification of the protrusion length of M. peptidophilum (n = 49 cells and 166 protrusions). **D,** Quantification of the protrusion growth rate (n = 31 protrusions). **E,** Representative trajectories of two migrating cells. **F/G,** TrackMate quantification of the confinement ratio (**F**) and the migration speed (**G**) (n = 21 cells from 3 cultures). **H,** Representative time lapse micrographs of M. peptidophilum with 0.5% DMSO (**top**) and 250ng/mL Swinholide A (**bottom**). Swinholide A arrests the rearrangement of the cell morphology. The corresponding experiments are shown in **Supp. Movie S12**.

### Gelsolins and profilins as putative regulators of Lokiactin polymerization

While eukaryotes have a repertoire of more than 150 actin binding proteins (ABP) available to fine tune actin polymerization, Promethearchaeota seem to rely mainly on gelsolins and profilins^22–24^. These were identified in a majority of the 15 closed and 396 meta-genomes (according to GTDB) of this group^4,37^ and were previously shown to regulate nucleation, bundling and severing of actin filaments in biochemical assays^22–24^. We combined proteomics and structural homology searches with FoldSeek^38^, as well as co-immunoprecipitation (Co-IP) with a Lokiactin antibody to find regulators of Lokiactin.

Our proteomic analysis confirmed the expression of gelsolins and profilins within the top 500 out of 3061 expressed proteins **(Fig. 5a, S4a)** and Co-IP experiments revealed several proteins associated with Lokiactin **(Fig. 5b, S4b)**. Among these, Lokiactin itself (log_2_ fold change of 4.44), an Arf GTPase, one profilin and five gelsolin-homology (GH) proteins were significantly enriched (log_2_ fold change > 1, p < 0.05). A second profilin and two BAR-domain proteins were also detected, though they did not reach statistical significance. A full list of all identified proteins is described in **Supplementary table 2.** It should be noted that, as is generally the case in proteomics, our Co-IP experiments are likely biased towards cytoplasmic proteins, leading to an underrepresentation of membrane-associated or transmembrane proteins.

**Figure 5:**
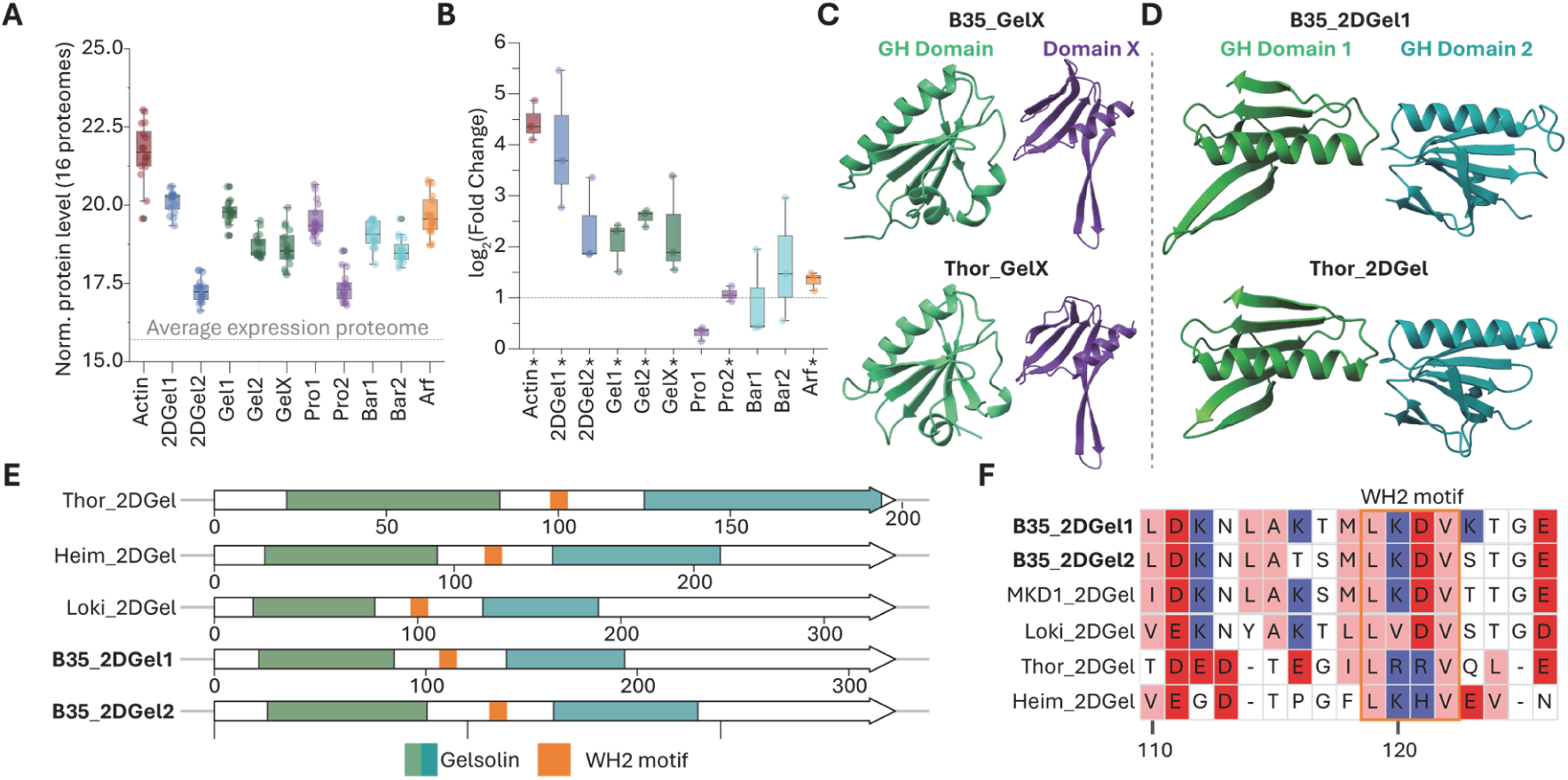
Co-immune precipitation and protein modelling A, A boxplot of the expression levels of selected proteins compared to the average expression level of all proteins from 16 proteomes (gray line). **B,** Box plot quantifying the fold change of the proteins in **(A)** in a Co-IP experiment using a Lokiactin antibody. Proteins with a (*) are enriched significantly (log2 fold change < 1, p < 0.01; Co-IP experiments were repeated with three cultures). **C,** Comparison of the AF3 modelled GH and disordered domain of unknown function (“X”) of B35_GelX and Thor_GelX. **D,** Comparison of the two AF3 modelled GH domains from B35_2DGel1 and Thor_2DGel. **E,** Comparison of the domain predictions of previously characterized 2D gelsolins and from L. ossiferum (in bold). **F,** Zoom-In showing the highly conserved WH2 motif of Promethearchaeota (LKDV) between the two GH domains. A list of NCBI identifiers of all proteins is in **Supplementary table 3**.

The detected profilin (B35_Pro2) shares high structural similarity with a previously biochemically characterized profilin from Odinarchaeota (RMSD_pruned_ = 0.896). This profilin was shown to slow actin polymerization, a mechanism which is reversible in the presence of phosphatidylinositol phosphate (PIP) lipids^22^ (**Fig. S4c**). We found homologues of three enzymes involved in PIP headgroup synthesis (myo-inositol-1-phosphate, phosphatidylinositol-phosphate synthase and Phosphoinositide 3-kinase), which are expressed within L. ossiferum (**Fig. S4d**).

Promethearchaeota gelsolins were previously shown to sever, cap, nucleate and bundle actin filaments in *in vitro* assays^23,24^. A phylogenetic analysis of L. ossiferum gelsolins clustered them into three distinct groups: B35_GelX contains two disordered domains and one GH domain and clustered with Thor_GelX, which was shown to disassemble actin filaments^23^ (**Fig 5c, S4e**).

Gelsolins with two GH domains (B35_2DGel) formed a sister group to eukaryotic villin and resembled the domain organization of other Promethearchaeota gelsolins, previously shown to sever or bundle actin filaments^23,24^ (**Fig. 5d, S4f**). These Promethearchaeota gelsolin homologues were described to contain WH2 motifs, which play a role in nucleating actin filaments and were shown to be sensitive to calcium signalling^24^. We confirmed the presence of this Asgard WH2 motif (LKDV) in both B35_2DGel proteins (**Fig. 5e, f**).

The remaining two gelsolins contain a single GH domain and one disordered domain and might represent an evolutionary transition state (B35_Gel), whose roles need further experimental validation (**Fig. S4g, Fig. S5**).

## Discussion

In this study we established anaerobic live-cell microscopy of L. ossiferum and M. peptidophilum, representing two different lineages of Promethearchaeota. We find these cells to dynamically rearrange their cell bodies and constantly form and retract protrusions, which are often branched. This plasticity is unprecedented in prokaryotes and usually associated with eukaryotic cells^39,40^. Since the membrane surface was kept constant, these shape changes do not seem to be facilitated by synthesis of novel lipids, uptake of lipids from syntrophic partners or externalisation of intracellular lipid compartments.

Furthermore, the cells display a unique type of motility, akin to amoeboid or crawling motion performed by immune cells or certain protists^41–43^. This motility is distinct from the hitherto described swimming motion in Archaea and Bacteria, which is driven by an archaellum or flagellum respectively^28^. Instead, L. ossiferum and M. peptidophilum use their dynamic protrusions to move while adhered to a surface, with a balance of adhesion strength and detachment being important for the migratory behaviour as our experiments with poly-L-lysin suggest.

Both the membrane rearrangements and the crawling motility involve the Lokiactin cytoskeleton, as we find several inhibitors of actin to arrest these dynamics. The various ranges of effects of different (eukaryotic) actin inhibitors may be caused by differences in protein sequence, the filament ultrastructure or perhaps impaired cell permeability.

Different from other types of surface motilities found in bacteria^44^, the motility in Promethearchaeota depends on an internal cytoskeleton. The observed enormous membrane plasticity and crawling motility could be beneficial in the natural environment of Promethearchaeota, the interstices of marine sediment^45–47^, as it might allow to scan their environment for nutrients or interaction partners or to facilitate more efficient nutrient uptake. Based on the similarities of the detected gelsolin and profilin homologues to those previously studied from other Promethearchaeota^22–24^, we imagine the Lokiactin cytoskeleton to be subject to dynamic regulation, possibly controlled by calcium signalling. However, we expect this regulation to be more intricate, as e.g. suggested by the presence of an ARF GTPase in our Co-IP experiments. Understanding Lokiactin polymerization and how it is affected by profilin, gelsolins and PIP lipids requires more biochemical investigations and will be subject of future work.

Our findings demonstrate that archaea likely evolved the ability to use actin for motility prior to the emergence of eukaryotes. Current understanding of actin based cell migration is heavily biased towards metazoans and amoebozoans with a more complex migration regulation machinery, including motor proteins, i.e. myosin and polymerases such as formins or Ena-VASP^48–53^. However, examples of motility in the absence of these proteins are known, as e.g. in red algae, but are studied in less detail^54–56^. Further investigations of the mechanism of Promethearchaeota motility should give mechanistic insights into an ancient form of actin-based cell migration.

The described plasticity and motility might have played an important role in eukaryogenesis, the emergence of eukaryotes, which remains one of the great enigmas in evolution. Current models include a merger of a Promethearchaeon with at least one bacterial partner^13,14,57–60^, in which the bacterium is eventually engulfed by the archaeon. Such a process necessitates plasticity of the archaeal membranes and cell shape, as shown by our study with L. ossiferum and M. peptidophilum. Furthermore, it is conceivable that the eukaryotic ancestor used protrusions to move towards potential interaction partners, connect to them to facilitate nutrient exchange and eventually engulf them.

Observing Promethearchaeota for an extended time in anaerobic conditions was a first step in understanding their dynamics and migration capacities. Future studies will aim at revealing the principles behind the observed behaviour and hopefully shed light on the events that happened about two billion years ago probably in deep marine sediments^61^ when the first eukaryotic cell arose.

## Methods

### Cultivation

*Ca.* L. ossiferum enrichments were grown protected from light in minimal lokiarchaeal media (MLM) in sealed serum bottles with a 80:20% N_2_:CO_2_ atmosphere (0.3 bars) at 20°C as described in^10^. *Ca.* M. peptidophilum HC1 was grown in casamino acids–peptone–yeast extract medium in sealed serum bottles with a 80:20% N_2_:CO_2_ atmosphere at 30°C as described in^12^.

### Imaging

Growth of L. ossiferum cultures was monitored by qPCR using Lokiarchaeal specific primers for 16S rRNA and was also used to select cultures for live cell imaging. Only cultures with more than 1*10_6_ L. ossiferum/mL and a relative abundance of L. ossiferum above 20% were chosen for experiments. At this stage the cells were either in late exponential or stationary phase and between 15 and 40 days old.

### Microscopy of fixed cells

For Immunostaining experiments culture was transferred on poly-L-lysin (0.01%) coated coverslips and incubated for 45 minutes in anaerobic conditions. Cells were then fixated with 3.7% PFA in PEM buffer (80 mM PIPES, 2 mM MgCl2, and 0.5 mM EGTA at pH 6.9). The cells were permeabilized with 0.1% Triton X-100 and blocked with 1% BSA solution in PBS-T. The primary antibody against Lokiactin and the secondary antibody + 1mg/mL Hoechst were diluted 1:1000 in PBS-T + 0.1% BSA and either incubated for 1h (primary) or 0.5 h (secondary) at room temperature, protected from light. In between these steps, coverslips were washed gently with PBS by changing the solution twice. Finally, the coverslips were mounted using the VectaShield antifading agent. To compare DNA signals and morphologies of L. ossiferum and their symbionts, the culture was fixed with 0.5% glutaraldehyde for 20 minutes. DNA was stained simultaneously with Hoechst 33342 (1:1000 diluted of 1mg/mL stock solution). After fixation, cells were centrifuged 2x for 10 minutes at 15000g and the resulting pellet was resuspended in milliQ to remove traces of Glutaraldehyde and media. The cell suspension was then dried onto slides and mounted using VectaShield antifading agent. Fixed cells were imaged with a Nikon Eclipse upright microscope equipped with a 100x Nikon 1.45 PlanApo oil objective.

### Preparation of flow chambers and live imaging

For upright phase contrast imaging, flow chambers were prepared with a Soda Lime microscopy slide (1*24*75 mm, bottom) and a Soda Lime coverslip (0.17*24*40mm, top). Two strips of double-sided sticky tape were placed on the outer edges of the bottom glass and sealed. After assembly, flow chambers were incubated in a 20:80% N_2_:CO_2_ atmosphere for at least 2 hours. 100µL of a Loki culture were sampled and transferred into an 2mL Eppendorf tube filled with 20:80% N_2_:CO_2_. Subsequently, 50µL of the culture were loaded into the flow chamber within an anoxic atmosphere. The flow chambers were sealed with Vaseline to avoid oxygen exposure. The cells were incubated for at least 90 minutes prior to starting imaging experiments. Timelapse experiments were performed with a Nikon Eclipse microscope, equipped with a 100x Nikon 1.45 PlanApo oil objective and a phase contrast setup. Movies were acquired with a frame rate of 2 seconds and for a maximum of 30 minutes.

For experiments with poly-L-lysine, the bottom slide was incubated with different concentrations of poly-L-lysine for 10 minutes, washed extensively with milliQ H_2_O and then dried for 2h at 60°C. Flow chambers were assembled and imaged as described above.

For inverted differential interference contrast (DIC) microscopy flow chambers were prepared as described above, but the bottom glass was exchanged with a Soda Lime coverslip (0.17*24*50mm). DIC experiments were performed with a Nikon Ti2-E microscope, equipped with a 100x Nikon 1.45 PlanApo oil objective. The flow chamber was mounted with a custom printed 3D slide holder into the imaging chamber. Experiments were recorded with frame rates of 5 - 60 seconds, determined by the number of channels imaged. During imaging, the temperature was kept at 22°C and a gas mix of 15%:85% N_2_:CO_2_ was flowed into the imaging chamber at a flow rate of 0.2mL/min to ensure an anoxic atmosphere. Oxygen levels were controlled by a Ibidi Gas Mixer and tracked with Anaerostrips in the experiment chamber.

For live imaging experiments with stained DNA, 1 drop of NucBlue was added to 1mL of culture prior to loading the flow chamber. Images were recorded every 60 seconds, with 20ms exposure of 1% 405nm laser. For live imaging of Lokiactin, FastAct was diluted 1:1000 directly into the culture prior to loading the flow chamber and incubated for at least 3 hours. Images were recorded every 10-30 seconds, using 50ms exposure time and 25% 640nm laser. For experiments with SiR-Actin and FastAct_X the same dilutions were used.

### Inhibitor experiments

Initial experiments of Actin inhibitors were performed at 25, 12.5 or 5µg/mL final concentrations (**Supplementary Table 1**). For titration experiments, the inhibitor stock solutions (5, 2.5 or 1 mg/mL) were diluted in steps of ten in DMSO and then diluted 1:200 directly with the Loki culture to titrate the inhibitor, while keeping DMSO constant at 0.5%.

### Image analysis

Image analysis was performed with FIJI^62^. Prior to color coding the protrusion growth, the movie was first stabilized targeting the cell body, smoothed with a walking average of 4 frames and then subsampled for every 30th frame. The resulting movie was colour coded using the temporal-Color code plugin and the mpl-magma lookup table.

To measure the rates of protrusion growth and shrinking, regions of interest with single L. ossiferum cells were selected from movies and stabilized using the cell body as a matching template. Then, a line was drawn along the protrusion of interest, and a kymograph was created using the reslice command. The rate was determined by measuring the travelled distance over time.

### Measuring protrusion dimensions and the change of length

To measure the length of protrusions over time, single cells were selected and the total length of all protrusions measured with the segmented line tool every 1 or 5 minutes. The change in length (ΔLength) was calculated by 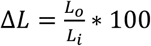 where L_0_ is the total length of the first time point and L_i_ the subsequent frames. The diameter of protrusions and the cell body of cells was determined from line profiles and the width of the resulting intensity profile at the full width half maximum.

### Tracking cell migration

To quantify cell migration we selected mobile cells and analysed regions of interest. Movies were inverted, their contrast normalized for analysis and cell bodies tracked with TrackMate^63^. Trajectories were built using the LAP tracker, allowing a linking max distance of 1µm, a gap-closing max distance of 2µm and a frame gap of maximum 5 frames. The resulting trajectories were manually curated, to avoid faulty linking due to focus shifts or floating of other, unbound cells into the FOV. Velocities, confinement ratio, Track durations and displacements were recovered from corresponding .csv and .xml files.

To determine the impact of protrusions on the migration direction, cells were manually segmented and used to train a pixel classification model in *ilastik* for automated detection of cell bodies and protrusion tips. Cell body trajectories were extracted with the *TrackMate* plug-in in Fiji. The movement direction was defined by the displacement vector between consecutive cell body centroids. For each frame, protrusion vectors were drawn from the cell centroid to each tip, and their alignment with the movement vector was calculated as the cosine of the included angle. Two metrics were derived: (i) the mean alignment, corresponding to the average cosine across all protrusions, and (ii) the weighted alignment, given by the sum of cosines weighted by protrusion lengths (tip-to-centroid distances) and normalized by total protrusion length. Both indices range from −1 (protrusions opposing movement) to +1 (protrusions aligned with movement).

### Effect of actin inhibitors

To assess the initial effect of inhibitors, all compounds were tested on the same culture within 3 subsequent days. We quantified the number of L. ossiferum cells with protrusion adhered to the glass slide after 90 minutes of incubation. For an in-depth analysis, the number of protrusions of each L. ossiferum cell in the presence of different inhibitors of Swinholide A were counted.

### Co-Immunoprecipitation

For fixation, 500 ml of cell culture were treated with formaldehyde to a final concentration of 0.25% for 20 min with gentle agitation. Crosslinking was quenched by the addition of glycine to a final concentration of 0.25 M for 5 min, followed by centrifugation and two washes with base medium. For immunoprecipitation, 10 µg of antibody was coupled using the Pierce Crosslink Magnetic IP/Co-IP Kit (Thermo Fisher), including crosslinking with DSS according to the manufacturer’s protocol. Cells were lysed in lysis buffer containing 50 mM Tris-HCl (pH 7.4), 150 mM NaCl, 10 mM MgCl₂, 1 mM EDTA, 0.1% IGEPAL CA-630, and a protease inhibitor tablet. Lysates were sonicated (1 s on/3 s off, 20% amplitude, 2 min) and centrifuged, and 200 µg of protein from the supernatant were incubated overnight with either antibody-coupled beads or control IgG-coupled beads (Abcam). Beads were washed twice with a detergent-free buffer (20 mM Tris-HCl pH 7.5, 75 mM NaCl).

### Sample preparation for mass spectrometry analysis

The beads were resuspended in 30 µL 2 M urea and 50 mM ammonium bicarbonate. The beads were digested with 150 ng LysC (mass spectrometry grade, FUJIFILM Wako chemicals) at room temperature for 90 minutes. The supernatant was transferred to a new tube. The beads were rinsed with 30µL 50mM ammonium bicarbonate and combined with the supernatant. Disulfide bonds were reduced with 2.4 µL of 250 mM dithiothreitol (DTT) for 30 min at room temperature before adding 2.4 µL of 500 mM iodoacetamide and incubating for 30 min at room temperature in the dark. The remaining iodoacetamide was quenched with 1.2 µL of 250 mM DTT for 10 min. Proteins were digested with 150 ng trypsin (Trypsin Gold, Promega) in 1.5 µL 50 mM ammonium bicarbonate overnight. The digest was stopped by the addition of trifluoroacetic acid (TFA) to a final concentration of 0.5%, and the peptides were desalted using C18 Stagetips^64^.

### Liquid chromatography-mass spectrometry analysis

LC-MS analysis was performed on a Vanquish Neo UHPLC system (Thermo Scientific) coupled to a timsTOF HT (Bruker). The system was equipped with a CaptiveSpray ion source (Bruker), and a column oven (Sonation).

Peptides were loaded onto a trap column (PepMap Neo C18 5mm × 300 µm, 5 μm particle size, Thermo Scientific) using 0.1% TFA as mobile phase, and separated on an analytical column Aurora Ultimate XT C18, 25 cm × 75 µm, 1.7 µm particle size, IonOpticks, applying a linear gradient starting with a mobile phase of 98% solvent A (0.1% FA) and 2% solvent B (80% acetonitrile, 0.08% FA), increasing to 35% solvent B over 60 min at a flow rate of 300 nl/min. The analytical column was heated to 50°C.

The mass spectrometer was operated in data-dependent acquisition (DDA) parallel accumulation serial fragmentation (PASEF) mode. MS2 data were acquired with ten PASEF scans per duty cycle. The ion mobility range was set to 0.64-1.42 V*s/cm, and the accumulation and ramp time was set to 100 ms. Precursor ions were selected for fragmentation using an isolation window of 2 m/z for m/z < 700 and 3 m/z for m/z > 700. The precursor intensity threshold was set to 2,500, precursor repetitions were enabled with a target intensity of 20,000, and active exclusion was enabled with a 0.4 min release delay.

Singly charged precursor ions were excluded using a polygonal filter applied in m/z-ion mobility space. TIMS elution voltages were calibrated linearly to obtain the reduced ion mobility coefficients (1/K0) using three Agilent ESI-L Tuning Mix ions (m/z 622, 922 and 1,222). Collision energy for fragmentation was scaled linearly with precursor mobility (1/K0), ranging from 20 eV (at 1/K0 = 0.6 V*s/cm) to 59 eV at (at 1/K0 = 1.6 V*s/cm).

### Mass Spectrometry data analysis

MS raw data were analysed with FragPipe (23.1), using MSFragger (4.3)^65^, IonQuant (1.11.12)^66^, and Philosopher (5.1.2)^67^. The default FragPipe workflow for label free quantification (LFQ-MBR) was used, except “Normalize intensity across runs” was turned off. Cleavage specificity was set to Trypsin/P, with two missed cleavages allowed. The protein FDR was set to 1%. Carbamidomethyl was used as fixed cysteine modification; methionine oxidation and protein N-terminal acetylation were specified as variable modifications. MS2 spectra were searched against the *Candidatus* L. ossiferum database with 5119 sequences, concatenated with a database of 379 common laboratory contaminants (release 2025_01, https://github.com/maxperutzlabs-ms/perutz-ms-contaminants).

Computational analysis was performed using Python and the in-house developed Python library MsReport (0.0.30) https://doi.org/10.5281/zenodo.15309090. Only non-contaminant proteins identified with a minimum of two peptides and being quantified in at least two replicates of one condition were considered for further analysis. LFQ protein intensities were log2-transformed and normalized across samples using the ModeNormalizer from MsReport. The ModeNormalizer method involves calculating log2 protein ratios for all pairs of samples and determining normalization factors based on the modes of all ratio distributions. Missing values were imputed by drawing random values from a normal distribution. Sigma and mu of this distribution were calculated per sample from the standard deviation and median of the observed log2 protein intensities (μ = median sample LFQ intensity – 1.8 standard deviations of the sample LFQ intensities, σ = 0.3 × standard deviation of the sample LFQ intensities). iBAQ intensities were calculated by dividing protein intensities by the number of theoretically observable tryptic peptides between 6 and 30 amino acids. To estimate the relative protein abundances within each sample, the iBAQ intensities were normalized by dividing them by the total sum of iBAQ intensities for all proteins in the sample. Statistical analysis was performed using the linear models for microarray analysis (limma) v.3.54.2 ^68^ package in R. Moderated *t*-statistics were calculated using the limma-trend method, and multiple testing correction was applied using the Benjamini–Hochberg method. The Python library XlsxReport (0.1.1, https://doi.org/10.5281/zenodo.15129818) was used to create formatted Excel files summarizing the results of the proteomics experiments.

### Gelsolin domain alignment and phylogenetic tree

Reference sequences for gelsolin-like homologs, including gelsolin, villins, and scinderin, were obtained from previous studies^23,69–71^ and retrieved from the National Center for Biotechnology Information (NCBI) database. The selected gelsolin sequences from L. ossiferum were aligned with these references using mafft v7.505 with the parameters “--auto” for multiple sequence alignment (MSA). Low-quality regions in the alignment were trimmed using TrimAl v1.4.rev15 with the parameters “-gt 0.95 -cons 50”, and a maximum-likehood tree was estimated with IQ-TREE2 v2.2.2.6 with 1000 ultrafast bootstraps. The best-fitting protein model, Q.pfam+F+R6, was determined with ModelFinder. The resulting tree was uploaded to iTOL for visualization.

Gene structure visualization was performed in R v4.5.0 using a combination of packages including ggplot2 for base graphics, gggenes for gene structure representation, and ggsci for scientific color palettes. Domain information was obtained with InterPro v106.0 database.

## Data availability

Plots were created using Jupyter Notebook. Source data for all figures have been provided in data source tables. Microscopy raw data for the supplementary videos are available from the first and corresponding author upon request ( and). The mass spectrometry proteomics data have been deposited to the ProteomeXchange Consortium via the PRIDE partner repository^72^ with the dataset identifier PXD068820.

## Acknowledgments

Proteomics analyses were performed by the Mass Spectrometry Facility at Max Perutz Labs using the VBCF instrument pool. This work would not have been possible without the technical support in cultivation of L. ossiferum from Karin Hager, Fatma Baraket, Andrea Malits and Sarah Harrer. We are thankful for the support by the Imaging and Optics Facility (IOF) at ISTA for setting up the anaerobic environment during imaging. We would also thank Philipp Klahn (University of Gothenburg, Sweden), Marc Stadler and members of his department (HZI Braunschweig) as well as Russell Cox (LUH Hannover) for providing different cytochalasans. This work was supported in part by the GermanResearch Council (DFG) through the CytoLabs consortium (Research Unit 5170 to K.R. and T.E.B.S.), by the Japan Society for the Promotion of Science (JSPS) Grant-in-Aid for Scientific Research 22H04985 to H.I. and M.K.N. Furthermore, R.H. was supported by CZI grant DAF2020-225401 and grant DOI https://doi.org/10.37921/120055ratwvi from the Chan Zuckerberg Initiative DAF. Finally this work was supported by the Austrian Science Fund (FWF) with W1257 and EFP 25 to C.S.

## Author contributions

P.R. and C.S. conceptualized experiments, P.R and T.V. performed and analysed microscopy experiments, R.H. performed morphodynamic analysis, N.M. and T.V. performed proteomic experiments and analysis, Z-H.L. performed bioinformatic analysis, K.S. characterized Actin inhibitors. P.R. and C.S. wrote the original draft. P.R., T.V., Z-H.L., N.M., K.S., M.N., S.B., T.E.B.S., K.R., H.I., M.S. and C.S. reviewed and edited the manuscript. C.S. and M.S. supervised the study.

---

**Supplementary Figure 1:**
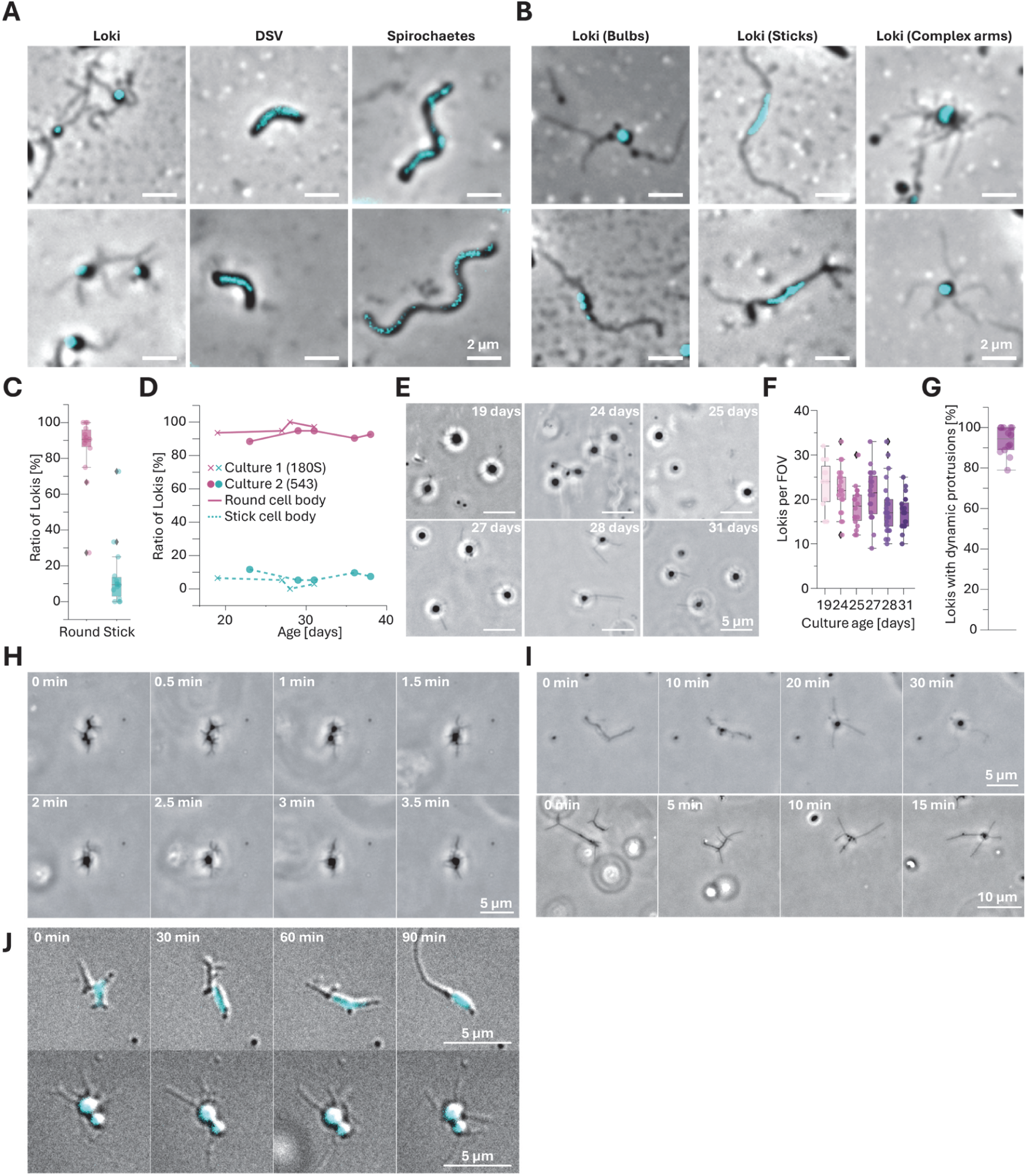
L. ossiferum undergo dynamic shape change on a minute time scale A, Representative micrographs of Glutaraldehyde fixed L. ossiferum and its syntrophic partners Desulfovibrio (DSV) and Spirochaetes stained with NucBlue. **B,** Representative micrographs of GA-fixed and NucBlue-stained L. ossiferum adopting different cell shapes**. C,** Ratio of round and stick shaped L. ossiferum cells in 9 cultures. **D,** Quantification of the round/stick ratio of two L. ossiferum cultures repeatedly imaged for 3 weeks**. E,** Representative micrographs of the same L. ossiferum culture at different time points. **F,** Quantification of the number of adhered L. ossiferum per FOV. **G,** Quantification of the percentage of L. ossiferum with dynamic protrusions. **H/I,** Representative PC time lapse micrographs of L. ossiferum undergoing dynamic shape changes. The corresponding experiments are shown in **Supp. Movie S4. J,** Representative DIC time lapse micrographs showing DNA-stained L. ossiferum either undergoing shape change (top) or retaining the same shape (bottom). The corresponding experiment is shown in **Supp. Movie S5.**

**Supplementary Figure 2:**
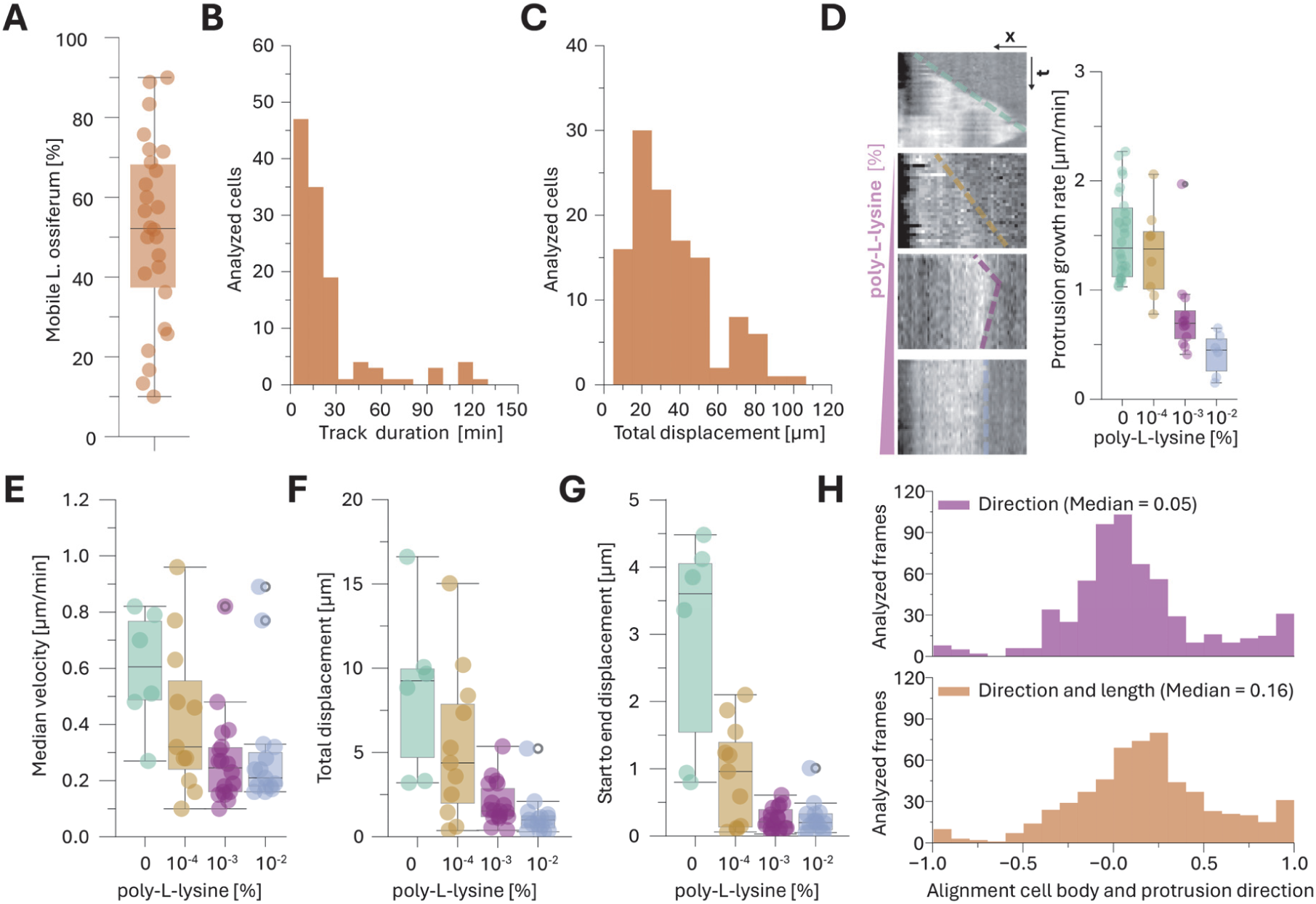
Poly-L-lysine and protrusion effect on L. ossiferum mobility A, Quantification of the percentage of mobile L. ossiferum per field of view from 12 different cultures. **B/C,** Quantification of the track duration **(B)** and total displacement **(C)** of motile L. ossiferum cells. **D,** Representative kymographs of protrusions of L. ossiferum adhered to glass with increasing concentrations of poly-L-lysine and the corresponding quantification. **E-G,** TrackMate quantification of L. ossiferum cells adhered to glass coated with increasing concentrations of poly-L-Lysine measuring the median instantaneous velocity **(E),** the total displacement **(F)** and the start to end displacement **(G)**. **H,** Histogram of the alignments of cell body displacement and the direction of the protrusions without (**top**) and with (**bottom**) weighting the protrusion length (n = 19 cells and 588 frames). Examples of the segmentation used for this analysis are shown in **Supp. Movie S8**.

**Supplementary Figure 3:**
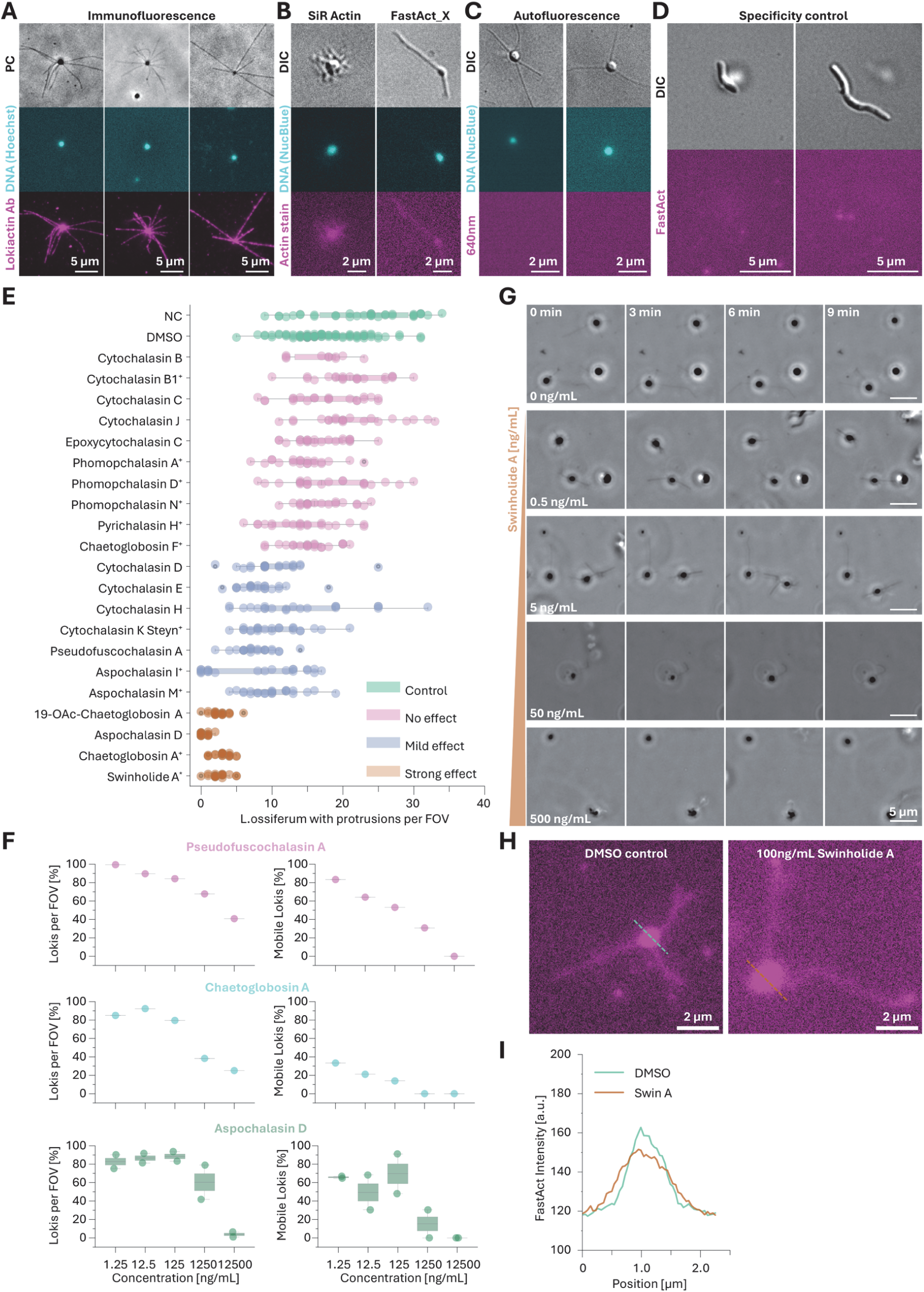
Controls for fluorescent imaging and actin inhibitor effect on L. ossiferum A, Representative micrographs of Immunofluorescence experiments of L. ossiferum stained with Hoechst (DNA, cyan) and an antibody targeting Lokiactin (magenta). **B,** Representative micrographs of L. ossiferum cells stained with NucBlue (DNA) and SiR-Actin and FastAct_X (Lokiactin). **C,** Autofluorescence controls of DNA stained L. ossiferum using the same imaging modalities as in experiments with FastAct. **D,** Specificity control of the FastAct stain. Halodesulfovibrio and Spirochaetes show no visible signal after incubation with FastAct for 24 hours. **E,** List of all tested inhibitors of eukaryotic Actin and their effect on the number of adherent L. ossiferum. The concentrations used were 25 (+), 12.5 and 5 (*) µg/mL and are listed in **Supplementary Table 1**. **F,** Titration experiments with three inhibitors of eukaryotic actin and a quantification of the average number of adhered L. ossiferum cells with protrusions (**left**) and their mobility (**right**) per experiment. **G,** Representative PC time lapse micrographs of L. ossiferum in the presence of increasing concentration of Swinholide A. The corresponding experiments are shown in **Supp. Movie S10**. **H**, Representative micrographs of FastAct-stained L. ossiferum in the absence (**left**) and presence (**right**) of 100ng/mL Swinholide A**. I,** Line profiles of the FastAct signal of the micrographs in **H**.

**Supplementary Figure 4:**
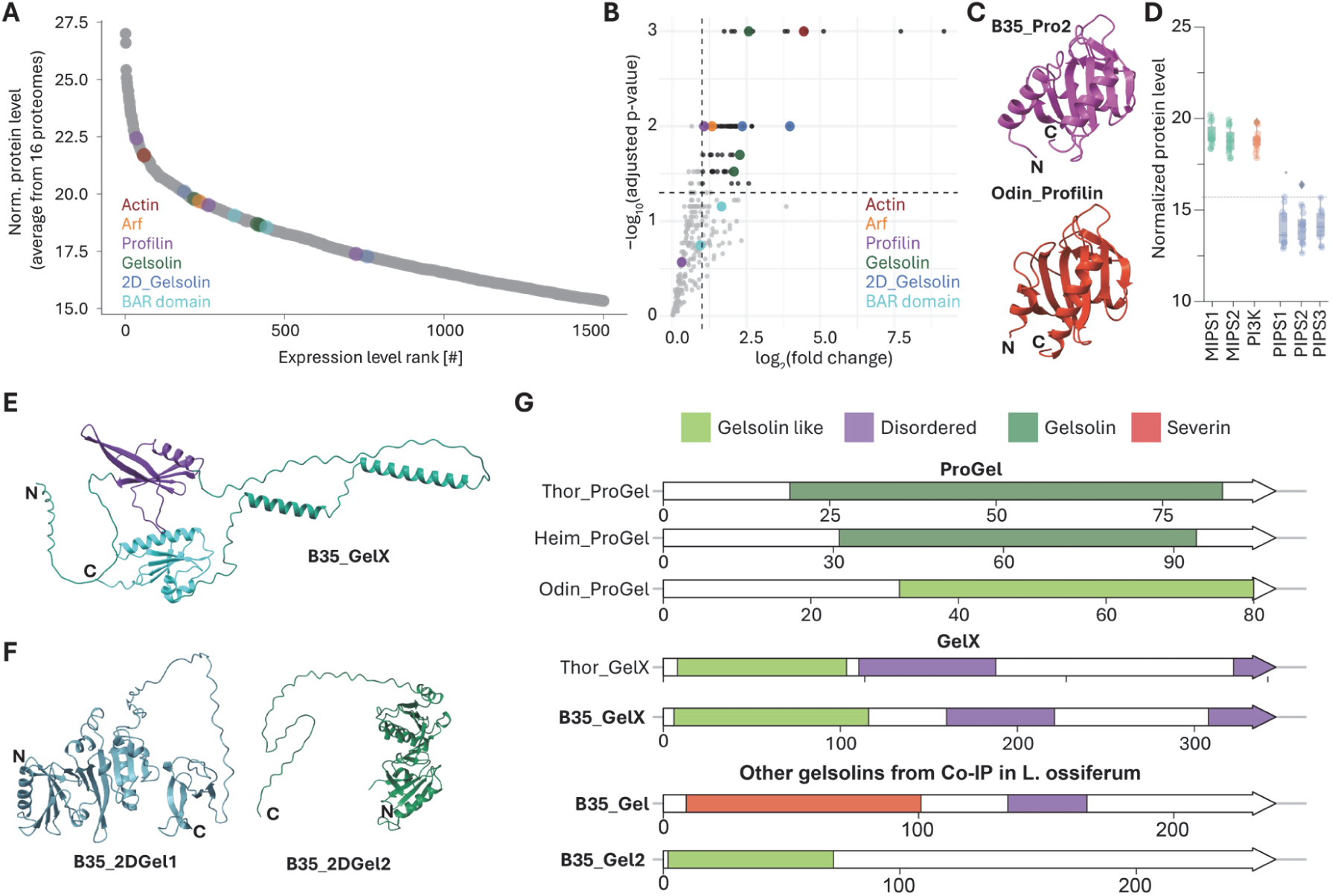
Protein expression, Co-IP and bioinformatic comparisons of gelsolins A, Expression plot of the top 1500 expressed proteins. Proteins identified in Co-IP are highlighted. **B,** Volcano Plot of proteins identified in a Co-IP experiment with actin antibodies. Proteins in the top right corner are enriched significantly. Coloured proteins are of particular interest for the regulation of actin filaments. **C,** AF3-predicted protein structures of L. ossiferum B35 profilin 2 and Odin profilin. **D,** Box plot of the expression levels of proteins involved in synthesis of PIP lipid headgroups. **E,** AF3-predicted structures of B35_GelX. Apart from the GH (green) and X domain (magenta), the protein contains an extensive linker of unknown function. **F,** AF3-predicted structures of L. ossiferum B35 gelsolins with two GH domains and their extensive C-terminal extensions of unknown function. **G,** Comparison of ProGel, a protein found in other Promethearchaeota but not in the Co-IP of L. ossiferum B35 followed by comparison of the domain structures of B35_GelX with Thor_GelX. Finally, the domains of the remaining gelsolins found in Co-IP experiments are shown. Their respective roles remain to be discovered

**Supplementary Figure 5:**
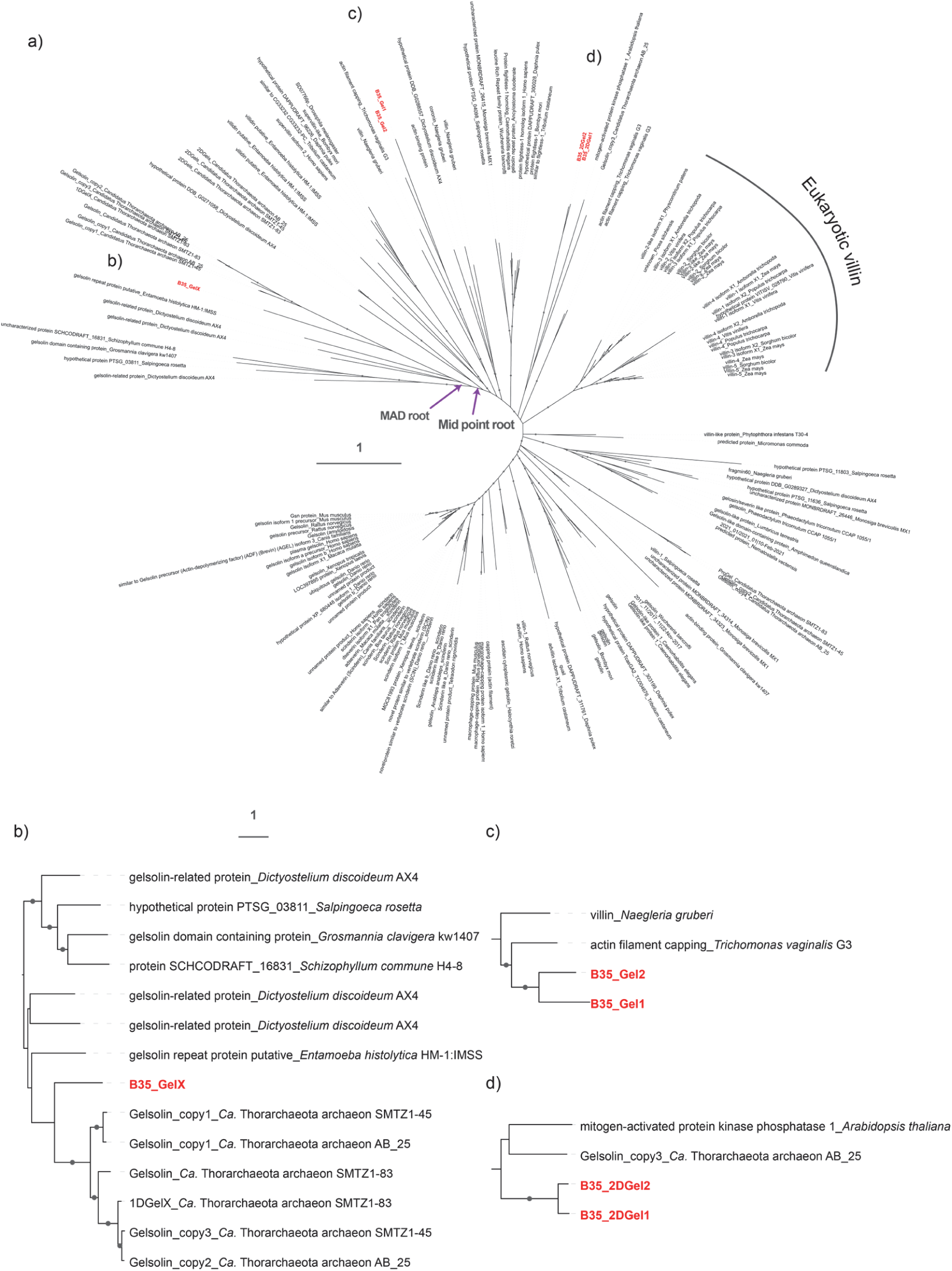
Maximum-likelihood phylogenetic tree of the homologs of gelsolin. Amino acid sequences from the tree were obtained from the NCBI database. a) The overview of all gelsolin homologs. Sequences from L. ossiferum B35 were marked in red. b-d) The enlarged phylogenies of gelsolin homologs from L. ossiferum B35. Nodes with ultrafast bootstrap value ≥ 80% (60%) were indicated as solid (hollow) circles.

